# LXA4 promotes the browning of white adipose through miR-133a-3p/SIRT1 pathway

**DOI:** 10.1101/2024.02.14.580287

**Authors:** Dan Yu, Yuan Ruan, Yisu Wang, Xiaopei Chen, Dan Wang, Tianfeng Wu

## Abstract

Lipoxin A4 (LXA4) promotes the browning of white adipose and energy consumption. The specific mechanism of which involved in white adipose browning is less clear. A high-fat diet (HFD) mouse model was constructed. Different groups of mice were treated with LXA4 accordingly. The body weight of mouse, subcutaneous and visceral fat, and food intake were recorded. The effect of LXA4 was examined by observing changes in pathology, serum insulin and lipid accumulation indices. The effects of LXA4/miR-133a-3p/Sirtuin1 on lipid droplet formation, fat browning-related genes, and the insulin receptor-AKT pathway in cells were examined after induction of adipocyte differentiation in 3T3-L1 precursors. At the cellular level, LXA4 promoted lipid droplet formation, expressions of fat browning genes and activation of the insulin receptor-AKT pathway in differentiated 3T3-L1 cells. MiR-133a-3p agomir partially offset the effects of LXA4. SIRT1 was a downstream target gene of miR-133a-3p, participating in the promotive effects of LXA4 on fat browning. LXA4 promotes white adipose browning and relieves insulin resistance through miR-133a-3p/SIRT1 pathway.

## Introduction

Obesity is a common metabolic syndrome developed when the human body ingests more calories than it consumes, with the excess calories stored in the body as a form of fat, which exceeds the normal physiological requirements (Mayoral et al., 2020). Epidemiological statistics indicated that obesity rate is increasing with each passing year, reaching 5% in children and 12% in adults as of 2015 (Afshin et al., 2017). The obesity rate appears to be lower in children than in adults, but the pertinent growth rate is even higher in children. Obesity and related metabolic diseases, such as type 2 diabetes, hypertension, cardiovascular and cerebrovascular diseases, tumors, etc. have become global public health problems (Erickson et al., 2015). Therefore, corresponding prevention and treatment are of positive significance.

The adipose tissue can be classified into white adipose tissue (WAT) and brown adipose tissue (BAT). WAT mainly functions in storing energy as a form of triacylglycerol in large lipid droplets, and secreting various endocrine hormones to balance the body’s energy (Montanari et al., 2017). The brown adipocytes are primarily feasible in oxidizing fat, dissipating heat, supplying energy, maintaining the body temperature balance, and preventing the body from hypothermia (Boon and van Marken Lichtenbelt, 2016). Under special conditions such as cold, some white adipocytes can be induced to become brown adipocytes (white adipocytes were interspersed with classic brown and beige adipocytes). Brown adipocytes can express uncoupling protein 1 (UCP1), thus producing UCP1-dependent heat, a process known as “browning” (Bargut et al., 2016, Ishibashi and Seale, 2010). The browning of white adipocytes enhances the oxidative decomposition of adipose tissues, promotes heat production, and reduces fat accumulation, accomplishing the goal of reducing fat and losing weight. Therefore, the promotion of BAT activation and WAT browning has emerged as a new orientation for the prevention and treatment of obesity and related metabolic diseases (Calvani et al., 2014).

Lipoxins, a kind of endogenous lipid mediators metabolized by Arachidonic acid, are produced by lipoxygenase, with main functions referring to the acceleration in timely resolution of inflammatory response, and obvious inhibition in driving and activation of white blood cells, which is an important anti-inflammatory mediator in the body (Hu et al., 2016, Romano et al., 2015). Lipoxin A4 (LXA4), as a vital member in this family, can inhibit inflammation, the generation of oxygen free radicals and the recruitment of neutrophils, as well as protect against oxidative stress-induced damage (Reis et al., 2017). Studies have proved that LXA4 reduces liver steatosis, partly because it promotes browning of WAT (Elias et al., 2016). However, how LXA4 participates in the process of WAT browning remains to be elucidated. This study used food-borne obesity mouse models and the induced differentiated adipocytes as the research subjects to probe into the specific underlying mechanism with regards to the regulation of LXA4 on the browning of WAT by detecting changes in lipogenesis, insulin, and adipocyte browning-related indicators.

## Materials and Methods

### Animal experiment

#### Feeding and obesity model construction

According to the relevant requirements of the Laboratory Animal Welfare Ethics Committee of Zhejiang Academy of Medical Sciences on animal protection (research license: 2019-131), 24 C57BL/6 mice (male; specific pathogen free (SPF) grade; Jiangsu ALF Biotechnology Co., LTD) were raised in a special animal experiment center and provided with an adequate diet, accompanied by the weight controlled within the range of 21 ± 2 g. The mice were given three days to acclimate to the breeding environment.

After the experiment officially started, the mice were randomly divided into 4 groups (6 mice/group) (Zheng et al., 2021): Control group, mice were fed with a normal diet for 9 weeks; Control + LXA4 group, mice were injected intraperitoneally with 5 ng/g/day LXA4 solution for 3 consecutive days, and then mice subsequently were fed with normal diet for 9 weeks.; High-Fat Diet (HFD) group, the mice were fed with a 60% HFD for 9 weeks; HFD + LXA4 group, the mice were injected intraperitoneally with 5 ng/g/day LXA4 solution for 3 consecutive days, and then mice were fed with a 60% HFD for 9 weeks. At the frequency of once a week, the weight of the mice was recorded on the same day, and the food intake of the mice was recorded after one week of rearing (1 time/week). LXA4 (GC18552) was provided by GLPBIO of the United States, while ordinary feed (SWS9112) and 60% high-fat feed (D12492) were provided by JIANGSU XIETONG PHARMACEUTICAL BIO-ENGINEERING CO., LTD.

#### Detection of obesity-related indicators

On the 64th day of the experiment, the mice were locally anesthetized around their eyes. The blood collected from the eye socket was used to detect the content of glucose (mg/dl), total cholesterol (TC; mg/dl), low-density lipoprotein cholesterol (LDL-C; mg/dl), high-density lipoprotein cholesterol (HDL-C; mg/dl), and alanine aminotransferase (ALT; U/L). The test kits used above were provided by J&L Biological (China): Mouse Glucose Test (JL18898), TC (JL18914), LDL-C (JL20313), HDL-C (JL20356) and ALT (JL17332).

#### Insulin sensitivity and glucose tolerance test

After the above-mentioned blood collection, all experimental mice were prohibited from eating for 12 hours (h) (allowed drinking water). The mice were injected (in the abdominal cavity) with 1.2 U/kg of insulin after fasting for 5 h, and 2 g/kg glucose solution (50%) orally after fasting for 12 h. Then their blood glucose levels (mg/dl) were measured at 0, 30, 60, 90, and 120 minutes (min) on both times.

#### Detection of insulin-related indicators (ELISA)

After the experiments mentioned above, the mice were anesthetized with sodium pentobarbital (injection dose 50 mg/kg; Sigma). The blood taken from the abdominal aorta was used to separate serum which was applied to detect the insulin (μg/L) and leptin levels (ng/ml) of the mice. The Homeostasis model assessment (HOMAIR) was performed based on the measured fasting blood glucose and insulin values (HOMAIR = fasting blood glucose × fasting insulin / 22.5). Insulin (E-EL-M2614) and leptin (E-EL-M3008) test kits were provided by Elabscience Company (China).

#### Collection of adipose samples

Since the blood was collected, the mice were quickly sacrificed. With reference to Chen’s operation method (Zheng et al., 2021), BAT, subcutaneous WAT (inguinal, armpit) and visceral WAT (epididymis, kidney, spleen) were separated, and the latter two were weighed. Hematoxylin-Eosin staining (HE) can visually display the pathological changes of adipocytes. Therefore, BAT and WAT were then immersed in 4% paraformaldehyde for fixation, and embedded in paraffin to make adipose slices. The deparaffinized slices were put into the HE Staining Kit (M027) produced by SHANGHAI GEFAN BIOTECHNOLOGY. CO. LTD. The slices of adipose tissue after mounting were observed for the pathological changes of BAT and WAT cells under the same field of view (100 times) through the OLYMPUS CX23 biological microscope.

#### Immunohistochemical staining

The dewaxed adipose slices obtained as described above were soaked in 3% H_2_O_2_ for 8 min to inactivate endogenous enzymes, followed by a 20-min treatment of antigen blocking. The above operations were completed at room temperature. Anti-SIRT1 antibody (ab189494, Abcam, USA) was diluted 400 times with the diluent and fully covered the adipose tissues. After an overnight incubation at 4 °C, the reaction reagent was replaced with Bio-Goat anti-mouse IgG working solution at 37 °C for 30 min. The last step was the DAB color rendering operation. Except for the Anti-SIRT1 antibody, all the reagents used in the experiment were provided by the mouse SP immunohistochemistry kit (SP0011, Solarbio, China). The mounted slices were placed under a microscope (100 times) to observe the positive expression.

#### Cell culture and intervention Ordinary culture conditions

The 3T3-L1 precursor adipocytes (CL-173™) provided by ATCC were inoculated in the complete medium (90% DMEM medium (30-2002™) and 10% Calf Bovine Serum (30-2030™) purchased from ATCC) recommended in the instructions, and incubated in a special incubator with stable conditions (37 °C, 5% CO_2_).

#### Inducing adipogenic differentiation

In order to better simulate the environment *in vivo*, we induced the adipogenic differentiation of 3T3-L1 cells. The 3T3-L1 cells that were normally cultured to the fifth/sixth passage were added to adipogenic differentiation induction medium which was prepared according to the formula provided by Claudia (Muscarà et al., 2019): DMEM medium, fetal bovine serum (10%), and MDI inducer (0.5 mM 3-isobutylmethylxanthine + 1 μM dexamethasone + 1 μg/ml insulin). The stimulation was continued for 2 days (Day 0), and the medium was again replaced with a DMEM maintenance medium containing only fetal bovine serum (10%) and insulin (1 μg/ml). The maintenance culture lasted for 8 days, during which the medium was changed every two days (Day 2, Day 4, and Day 6). The control cells were always maintained in complete culture medium.

#### Grouping and LXA4 intervention

According to different intervention conditions, the cells were divided into Control group, MDI group, MDI + LXA4 (0.1) group, MDI + LXA4 (1) group, MDI + LXA4 (10) group and MDI + LXA4 (100) group. While MDI inducer or DMEM maintaining the medium intervention (Day 0, Day 2, Day 4, and Day 6), different concentrations of LXA4 (0.01, 0.1, 1, and 10 nM) were used to stimulate 3T3-L1 cells in the corresponding group. On Day 9, 3T3-L1 cells in each group were collected. In order to better observe lipogenesis, the microscope field of view was magnified 400 times. The lipid structure formed in the cell was identified and marked (red arrow).

#### Overexpression treatment of miR-133a-3p

MiR-133a-3p agomir (agomiR-133a-3p; miR40000145-4-5) and its negative control (agomir-NC; miR4N0000001-4-5) provided by Guangzhou RIBOBIO were transfected into MDI-treated 3T3-L1 cells through liposomes (by riboFECT CP Transfection Kit; C10511-05). 24 h latter, the transfection efficiency was measured through Quantitative Real-Time Polymerase Chain Reaction (qRT-PCR). The following are the experimental groups:3T3-L1 cells in control group were induced by MDI; 3T3-L1 cells in LXA4 (10) group were induced by MDI induced and 10 nM LXA4; 3T3-L1 cells in LXA4 (10) + agomiR-NC group were induced by MDI, 10 nM LXA4 and agomiR-NC; 3T3-L1 cells in LXA4 (10) + agomiR-133a-3p group were induced by MDI, 10 nM LXA4 and agomiR-133a-3p.

#### Sirtuin1 (SIRT1) silence/overexpression treatment

According to the targeting sequences as listed below, GenePharma (China) synthesized the plasmid of short hairpin RNA targeting SIRT1 (shSIRT1) for this study by: shSIRT1-1, 5’-CGCCTTTAGTGAAGTTATAGTTT-3’; and shSIRT1-2, 5’-CGCAACATTTTTAGATTAACAAA-3’. SIRT1 overexpression plasmid was obtained by amplifying the full length of SIRT1 and inserting into the pcDNA3.1 (-) overexpression vector (V795-20, Invitrogen, USA). ShSIRT1, SIRT1 overexpression plasmid and their respective negative controls (shNC; NC) were transfected into 3T3-L1 cells treated with MDI differentiation according to the aforementioned transfection method. All 3T3-L1 cells in this part of the experimental groups have undergone MDI differentiation treatment: MDI differentiated 3T3-L1 cells in LXA4 (10) group were treated by 10 nM LXA4; MDI differentiated 3T3-L1 cells in LXA4 (10) + shNC group were transfected with shNC and treated by10 nM LXA4; MDI differentiated 3T3-L1 cells in LXA4 (10) + shSIRT1 group were transfected with shSIRT1 and treated by 10 nM LXA4; MDI differentiated 3T3-L1 cells in LXA4 (10) + agomiR-133a-3p group were treated transfected with agomiR-133a-3p and by 10 nM LXA4;MDI differentiated 3T3-L1 cells in LXA4 (10) + agomiR-133a-3p + NC group were transfected with agomiR-133a-3p and NC, and treated by 10 nM LXA4; MDI differentiated 3T3-L1 cells in LXA4 (10) + agomiR-133a-3p + SIRT1 group were transfected with agomiR-133a-3p and SIRT1, and treated by 10 nM LXA4.

#### Oil Red O staining

The cell-specific Oil Red O staining solution (G1262) developed by Solarbio (China) can absorb smaller lipid droplets, which, therefore, was used to identify the lipogenesis in 3T3-L1 cells. The collected LXA4-intervented cells were first fixed with Oil Red O fixative for 30 min to reduce cell loss. After immersion in isopropanol, the newly prepared Oil Red O staining solution was added to the cells to be tested and cells were immersed for 15 min. After washing away the excess solution, the dyeing result was observed through a microscope (200 times), with staining from orange to red as a positive result.

#### QRT-PCR

The total RNA in 3T3-L1 cells or mouse WAT was extracted with TRIzol (15596018) produced by Invitrogen. Since the detection of mRNA and miRNA was designed in the experiment, the TaKaRa Reverse Transcription Kit (RR047A) for mRNA and the HaiGene TaqMan miRNA cDNA Synthesis Kit (D1802) for miRNA were selected for the next cDNA synthesis (Mohammadi-Yeganeh et al., 2013). The collected cDNA followed the same operation to complete the following qRT-PCR with SYBR Green (04913914001, Roche, Sweden) as the dye. After completing the test through Thermal Cycler-T100 gradient PCR machine (Bio-Rad, USA), we calculated 2^-ΔΔCt^ (the final form of the relative level of mRNA) based on the measured Ct value. During the experiment, β-actin (for mRNAs) and U6 (for miRNAs) played the roles of housekeeping genes. Due to the large number of detected genes, we listed the primer sequences synthesized by Shanghai Sangon Company in Table 2.

### Western blot

Similar to qRT-PCR, to complete the western blot experiment of gene protein level analysis, we first extracted the total protein in 3T3-L1 cells or mouse WAT. Thermo Scientific™ RIPA Lysis Buffer (89901) and Pierce™ Rapid Gold BCA Protein Assay Kit (A53227) were used for the extraction of protein and the determination of concentration, respectively, providing a guarantee for subsequent testing. The subsequent protein isolation process was completed by SDS-PAGE electrophoresis and electroporation. Immobilon-PSQ polyvinylidene fluoride (PVDF) membrane (0.2 µm; ISEQ00010, Millipore, USA) was used as a protein carrier in this step. The fully sealed PVDF membrane was soaked in the diluted primary antibodies overnight (4 °C), and then transferred to secondary antibodies to react for 1.5 h (room temperature). The excess secondary antibody diluent was washed away, the PVDF membrane was transferred to the Gel Doc™ XR+ imaging system (Bio-Rad, USA), and Super ECL Detection Reagent (36208ES60, YEASEN, China) was added dropwise to help the protein develop color. The relative protein level was finally expressed in the form of target protein/β-actin. The following antibodies were used in the experiment: phospho(p)-IR tyr972 (1/800, 155 kDa; 07-838, Sigma-Aldrich, USA), p-IRS1 r307 (1/800, 185 kDa; 05-1087, Sigma-Aldrich, USA), p-AKT (1/800, 56 kDa; ab38449, Abcam, USA), AKT (1/500, 56 kDa; ab8805, Abcam, USA), SIRT1 (1/1000, 80 kDa, ab189494, Abcam, USA), β-actin (1/4000, 42 kDa; ab8226, Abcam, USA), Rabbit Anti-Mouse antibody (1/8000, ab6728, Abcam, USA), and Goat Anti-Rabbit antibody (1/8000, ab6721, Abcam, USA). Prediction and verification of downstream target genes The TargetScan website (http://www.targetscan.org/vert_72/) was used to predict the binding sites of SIRT1 and miR-133a-3p, and the dual luciferase reporter assay was hereby designed to verify the results. The wild-type and mutant-type sequences of SIRT1 (SIRT1-WT, GGAACAGCUAAUCUAGACCAAAG, SIRT1-MUT, GGAACAGCUAAUCUAGUGCUAUG) were inserted into the pmirGLO vector (VT1439, YouBio, China). The obtained reporter genes were co-transfected with agomiR-133a-3p (or agomir-NC) into conventionally treated 3T3-L1 cells. The subsequent detection of luciferase activity was completed with the participation of the dual luciferase reporter gene detection kit (D0010-100T, Solarbio, China) and the Promega GLOMAX analyzer.

### RNA immunoprecipitation (RIP)

The RNA Immunoprecipitation Kit (P0102) obtained from GENESEED can more accurately verify the binding of miR-133a-3p and SIRT1. Specifically, it included the following three steps: magnetic beads and cells were separately pretreated; then IgG antibody and Ago2 antibody (ab186733, Abcam, USA) provided by the kit were used for the connection of antibody and magnetic beads; RNA was finally extracted from the collected complex with its level detected by PCR.

### Statistical Analysis

The statistical analysis of experimental data was completed by GraphPad Prism 8.0 software (USA). The contents of obesity-related indicators in Table 1 (glucose, TC, LDL-C, HDL-C, and ALT) were all expressed in the form of mean ± standard deviation. The analysis involved the difference comparison of multiple grouped data. Here, the one-way ANOVA and Bonferroni between-group analysis methods were unified. When *p*<0.05, the difference was considered to be statistically significant.

#### Result

### LXA4 treatment alleviated the increase in body weight and fat mass of mice caused by HFD

From the body weight data of figure 1A and the food intake records of figure 1B, it could be concluded that the HFD significantly increased the weight of the mice, with gradually decreased food intake (*p*<0.001, figure 1A-B). LXA4 treatment notably reduced the body weight of mice with HFD (*p*<0.001, figure 1A), but exerted few effects on food intake (figure 1B) of mice with HFD and the body weight and food intake of normally reared mice (figure 1A-B). Besides, HFD and LXA4 treatment also greatly affected the fat weight of mice. It can be clearly seen in the digital photos and histograms of figure 1C that the weight of subcutaneous and visceral fat in the mice in HFD group augmented dramatically. The treatment of LXA4 reduced the fat weight of obese mice (*p*<0.001, figure 1C). HE staining revealed the changes of WAT and BAT in mice treated with HFD and LXA4. The results showed that the cell volume in mice of WAT of HFD group rose markedly, while that in BAT reduced significantly (figure 1D). The above pathological changes were remarkably improved after LXA4 treatment.

**Figure 1.**
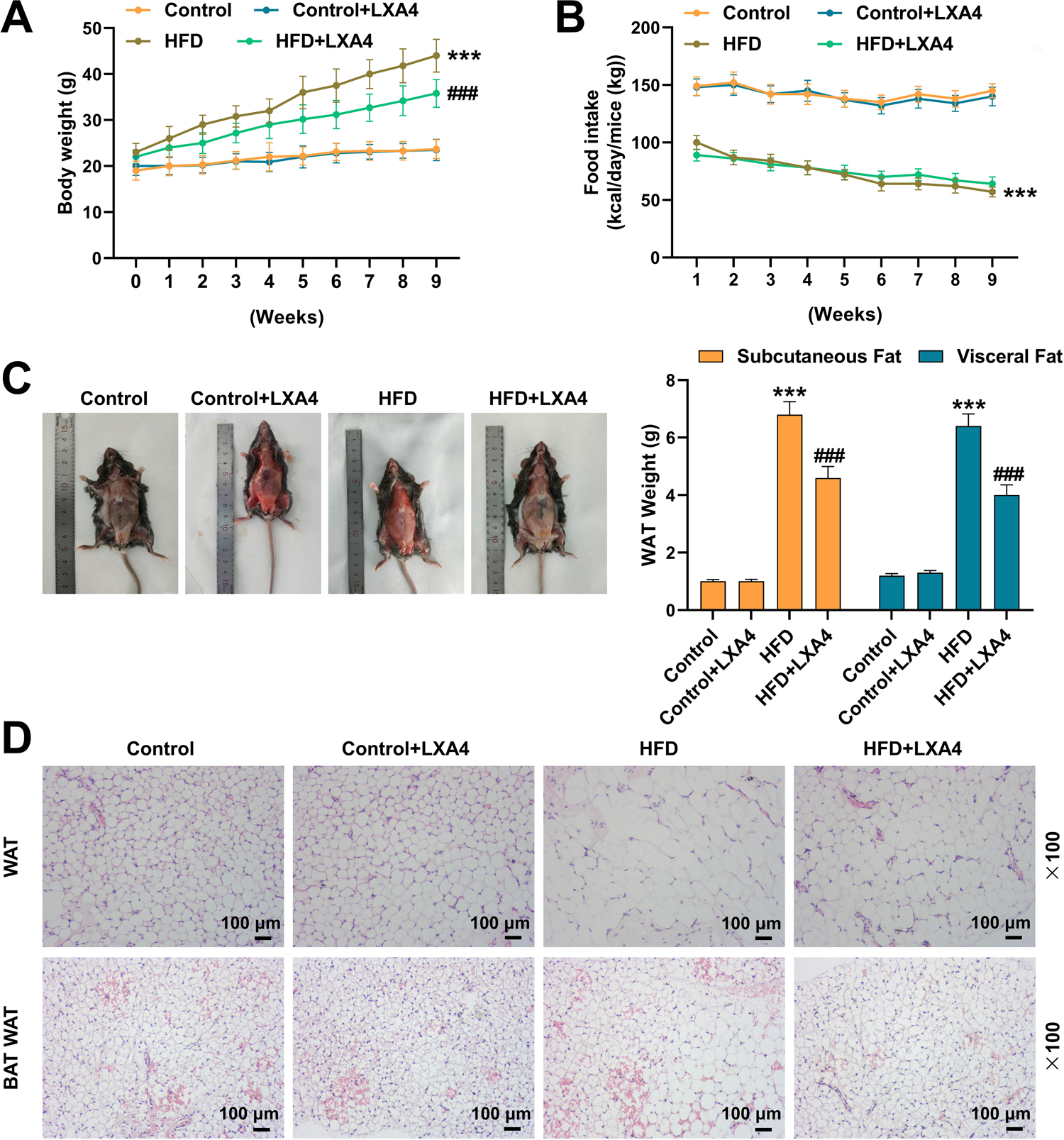
LXA4 treatment alleviated the increase in body weight and fat mass of mice caused by HFD. (A-B) The effects of High-Fat Diet (HFD) and Lipoxin A4 (LXA4) treatments on the body weight and food intake of mice were recorded. (C) The effects of HFD and LXA4 treatments on subcutaneous and visceral fat in mice were measured and analyzed. (D) The effects of HFD and LXA4 treatments on white adipose tissue (WAT) and brown adipose tissue (BAT) of mice were observed by Hematoxylin-Eosin staining. The magnification was 100 times. All experiments were repeated three times to average. \*\*\**p*<0.001 vs Control; ^###^*p*<0.001 vs HFD. Animal groups (n=6): Control group, Control + LXA4 group, HFD group, HFD + LXA4 group. The intraperitoneal injection dose of LXA4 was 5 ng/g/d.

### LXA4 treatment mitigated the abnormal changes in lipid accumulation and insulin sensitivity-related indicators caused by HFD

We analyzed the content of glucose and lipid accumulation-related indicators in the serum of mice. As depicted in Table 1, except for the ALT content, the levels of Glucose, Total Cholesterol, LDL-C, HDL-C and Triglyceride in the serum of mice in the HFD group mice were overtly lower than those of the control group (*p*<0.05, Table 1). After LXA4 treatment, the above-mentioned reduced indexes in the serum all appeared to be up-regulated to varying degrees, while the content of ALT in the serum of mice was downregulated. Figure 2 described the changes in insulin-related indicators. We can clearly note that HFD treatment resulted in a large accumulation of insulin and leptin in mice, with higher insulin sensitivity, glucose tolerance and HOMA-IR levels when compared with those in normal control mice (*p*<0.001, figure 2A-G). Similar to the previous one, the therapeutic effect of LXA4 was mirrored in the down-regulation of the above abnormally elevated indicators (*p*<0.001, figure 2A-G).

**Figure 2.**
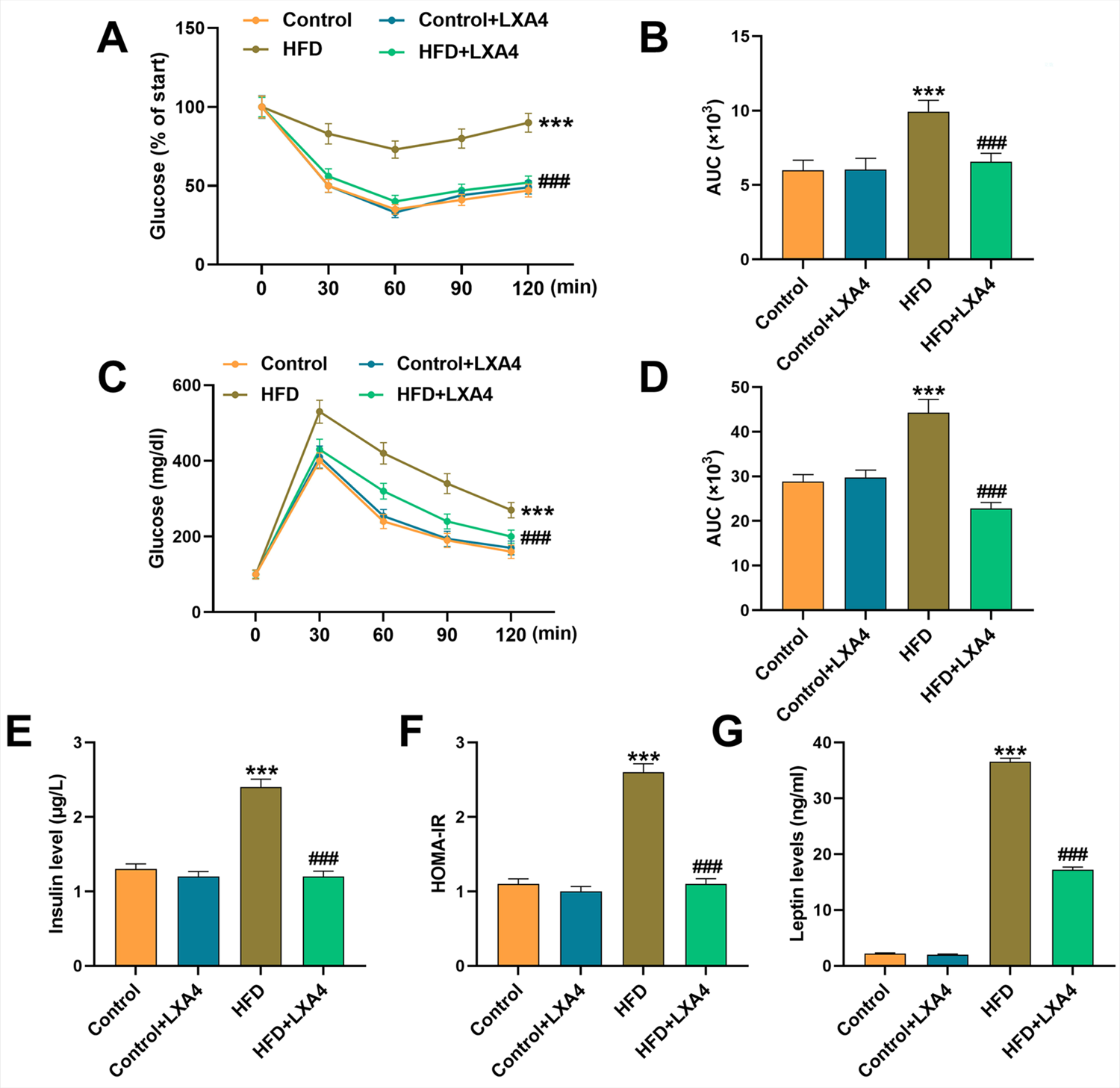
LXA4 treatment alleviated the abnormal changes in lipid accumulation and insulin sensitivity-related indicators caused by HFD. (A-D) LXA4 treatment alleviated the increase in insulin sensitivity and glucose tolerance of mice induced by HFD treatment. (E-G) The effects of HFD and LXA4 treatment on the levels of insulin and leptin in the serum of mice were tested by ELISA. HOMAIR = fasting blood glucose × fasting insulin / 22.5. All experiments were repeated three times to average. \*\*\**p*<0.001 vs Control; ^###^ *p*<0.001 vs HFD. Animal groups (n=6): Control group, Control + LXA4 group, HFD group, HFD + LXA4 group. The intraperitoneal injection dose of LXA4 was 5 ng/g/d.

### LXA4 promoted the browning of 3T3-L1 adipocytes and prevented the up-regulation of miR-133a-3p

Figures 3A and 3B delineated the effects of MDI and LXA4 induction on the formation of lipid droplets in 3T3-L1 cells. The red arrow in figure 3A and the red positive expression in figure 3B both marked the formation of lipid droplets. The number of lipid droplets in 3T3-L1 cells was abundant after MDI induction, which changed a little after the intervention of LXA4 at 0.01 nM or 0.1 nM, but evidently increased when the concentration of LXA4 reached 1 nM and 10 nM. Thus, 1 nM and 10 nM LXA4 were singled out for subsequent experiments. Next, we analyzed the expressions of fat browning-related genes in each group of cells. MDI induced the up-regulation of uncoupling protein 1 (UCP1), PR domain-containing 16 (PRDM16), peroxisome proliferator activated receptor coactivator-1α (PGC1-α), cytochrome c (Cytc) and cytochrome c oxidase subunit 4b (COX4b) (*p*<0.001, figure 3C), which were further strengthened after increasing the concentration of LXA4 (*p*<0.001, figure 3C), confirming that LXA4 can promote the browning of adipocytes and activate mitochondrial function.

**Figure 3.**
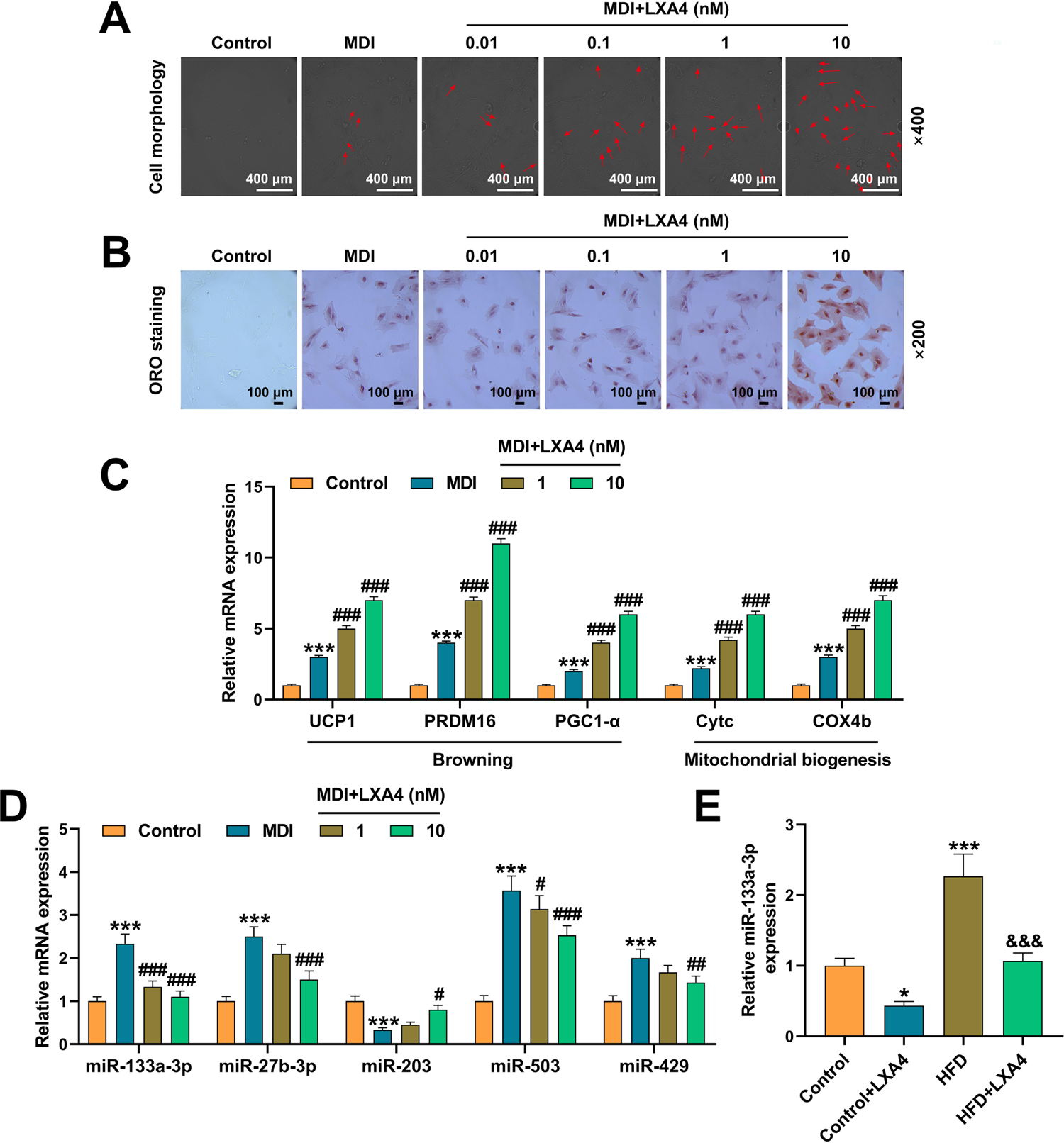
LXA4 promoted the browning of 3T3-L1 adipocytes and prevented the up-regulation of miR-133a-3p. (A) The effect of LXA4 on the morphology of 3T3-L1 adipocytes induced by MDI was observed under a microscope. The magnification was 400 times, and the red arrow marked the position of the lipid droplets. (B) The effect of LXA4 on MDI-induced lipid droplet formation in 3T3-L1 adipocytes was observed by oil red O staining. The magnification was 200 times. (C) The effect of LXA4 on the expression of adipocyte browning-related genes in 3T3-L1 cells induced by MDI was tested by qRT-PCR. β-actin acted as the housekeeping gene. (D) The effects of LXA4 on the expression of adipocyte browning-related miRNAs in 3T3-L1 cells induced by MDI were detected by qRT-PCR. U6 acted as the housekeeping gene. (E) The effect of LXA4 on the expression of miR-133a-3p in WAT was tested by qRT-PCR. U6 acted as the housekeeping gene. All experiments were repeated three times to average. \**p*<0.05, \*\*\**p*<0.001 vs Control; ^#^*p*<0.05, ^##^*p*<0.01, ^###^*p*<0.001 vs MDI; ^&&&^*p*<0.001 vs HFD. The concentration of LXA4 treatment was 0.01, 0.1, 1 and 10 nM, respectively.

Based on the important regulatory role of miRNAs in white adipose browning, we selected miRNAs related to white adipose browning in existing studies (Kim et al., 2019, Petrek and Yu, 2019, Guo et al., 2019, Man et al., 2020, Ye et al., 2021), and analyzed the changes of miRNAs under the treatment of LXA4. Compared with the MDI induction group, 10 nM LXA4 notably down-regulated the expressions of miR-133a-3p, miR-27b-3p, miR-503 and miR-429, and up-regulated that of miR-203 (*p*<0.05, figure 3D). Here, miR-133a-3p, with apparent changes, was selected to continue our research. The same results were also verified in mouse WAT. Specifically, an up-regulated expression miR-133a-3p were evidenced in mice subjected to HFD, which was prevented by LXA4 treatment (*p*<0.001, figure 3E).

### Up-regulation of miR-133a-3p offset the fat-browning effect of LXA4

AgomiR-133a-3p was transfected into MDI-induced 3T3-L1 adipocytes to analyze the role of miR-133a-3p in the promotive effects of LXA4 on fat browning. LXA4 treatment down-regulated the expression of miR-133a-3p in MDI-induced cells, which was effectively reversed by the transfection of agomiR-133a-3p (*p*<0.001, figure 4A). In the following lipid droplet formation and fat-browning gene expression tests, agomiR-133a-3p effectually inhibited LXA4-stimulated lipid droplet formation and the activation of fat-browning genes (UCP1, PRDM16, PGC1-α, Cytc, and COX4b) (*p*<0.001, figure 4B-D). These results confirmed our conjecture that LXA4 regulates miR-133a-3p and promotes adipocyte browning. Hence, we continued to analyze the protein changes of the insulin receptor-AKT pathway in the cells, proving that LXA4 treatment up-regulated p-IR tyr972, p-IRS1 r307, and p-AKT expressions, with activated insulin receptor-AKT pathway (*p*<0.001, figure 4E), while the superimposed intervention of agomiR-133a-3p partially reversed such effect on the activation of pathway (*p*<0.001, figure 4E).

**Figure 4.**
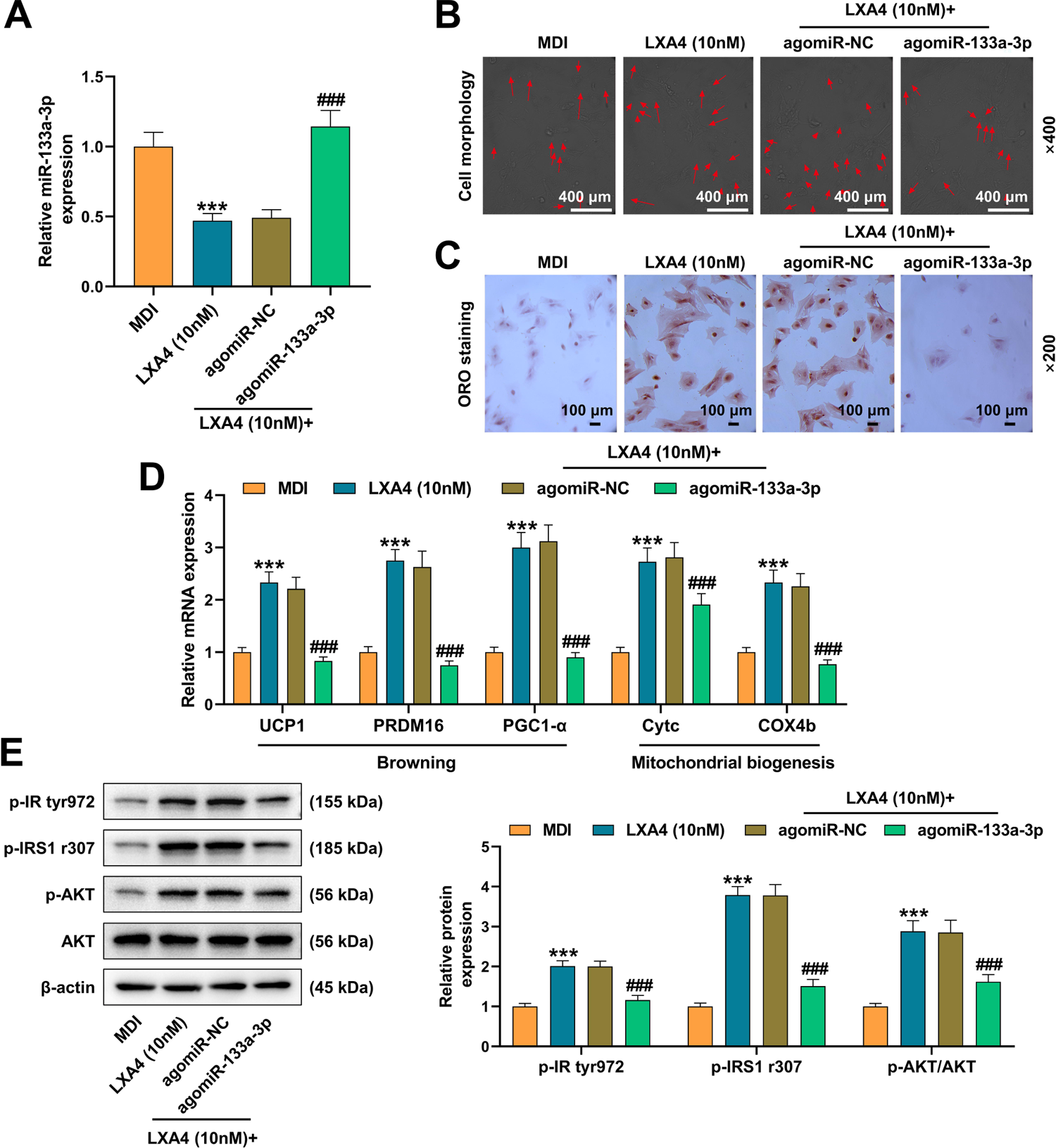
Up-regulation of miR-133a-3p counteracted the fat-browning promotion effect of LXA4. (A) The effects of LXA4 and miR-133a-3p agomir (agomiR-133a-3p) on the expression of miR-133a-3p in 3T3-L1 cells were measured by qRT-PCR. U6 acted as the housekeeping gene. (B) The effects of LXA4 and agomiR-133a-3p on the morphology of 3T3-L1 cells induced by MDI were observed under a microscope. The magnification was 400 times, and the red arrow marked the position of the lipid droplets. (C) The effects of LXA4 and agomiR-133a-3p on lipid droplet formation in 3T3-L1 cells induced by MDI were observed by oil red O staining. The magnification was 200 times. (D) The effects of LXA4 and agomiR-133a-3p on the expression of adipocyte browning-related genes in MDI-induced 3T3-L1 cells were detected by qRT-PCR. β-actin acted as the housekeeping gene. (E) The regulation of LXA4 and agomiR-133a-3p on the insulin receptor-AKT pathway in MDI-induced 3T3-L1 cells was tested by western blot. β-actin acted as the housekeeping gene. All experiments were repeated three times to average. \*\*\**p*<0.001 vs MDI; ^###^ *p*<0.001 vs LXA4 (10nM) + agomiR-NC.

### SIRT1 was a downstream target gene of miR-133a-3p, and its expression was regulated by LXA4 and agomiR-133a-3p

Figure 5A profiled the clear binding sites between SIRT1 and miR-133a-3p. The dual luciferase reporter assay and RIP in figure 5B and 5C fully proved the binding of SIRT1 and miR-133a-3p (*p*<0.001, figure 5B-C). The regulation of SIRT1 expression by LXA4 and miR-133a-3p was tested in various ways: In WAT of mice with HFD, LXA4 treatment significantly up-regulated the mRNA and protein expressions of SIRT1 (*p*<0.001, figure 5D, 5E and 5H), whilst in MDI-induced 3T3-L1 cells, LXA4 also promoted the expression of SIRT1, but agomiR-133a-3p offset the regulatory effect of LXA4 and down-regulated SIRT1 expression (*p*<0.05, figure 5F-G).

**Figure 5.**
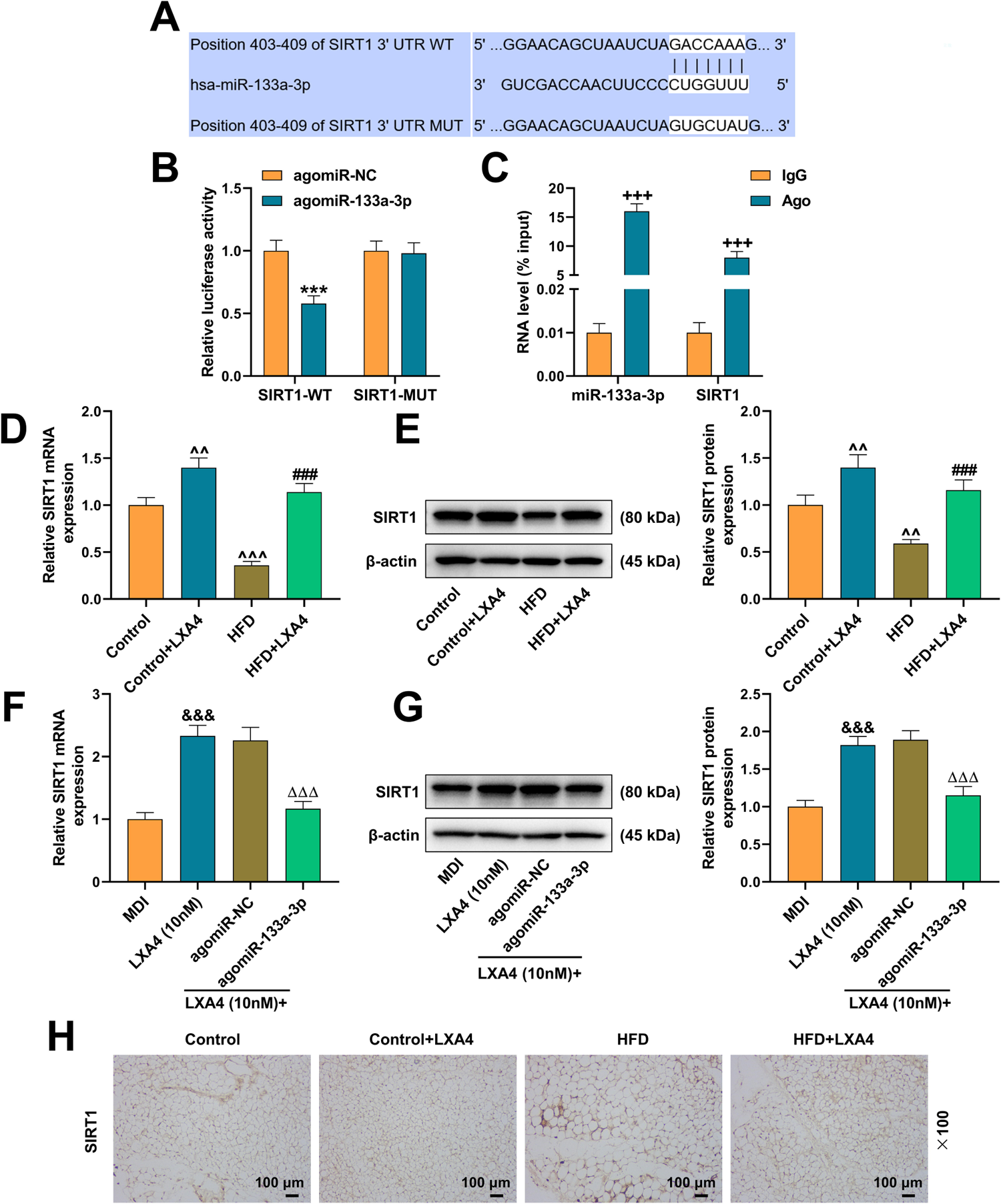
SIRT1 was a downstream target gene of miR-133a-3p, and its expression was regulated by LXA4 and agomiR-133a-3p. (A-B) The binding sites of SIRT1 and miR-133a-3p were predicted (TargetScan website) and verified (dual luciferase experiment). (C) The binding of SIRT1 and miR-133a-3p was verified by RNA immunoprecipitation (RIP) experiment. (D-E) The regulation of LXA4 on the expression of SIRT1 in mouse WAT was analyzed by qRT-PCR and western blot. β-actin acted as the housekeeping gene. (F-G) The regulation of LXA4 and agomiR-133a-3p on the expression of SIRT1 in MDI-induced 3T3-L1 cells was analyzed by qRT-PCR and western blot. β-actin acted as the housekeeping gene. (H) The positive expression of SIRT1 in mouse WAT was observed by immunohistochemical staining experiments. The magnification was 100 times. All experiments were repeated three times to average. \*\*\**p*<0.001 vs agomiR-NC; +++*p*<0.001 vs IgG; ^^*p*<0.01, ^^^*p*<0.001 vs Control; ### *p*<0.001 vs HFD; &&*p*<0.01, &&&*p*<0.001 vs MDI; △*p*<0.05, △△△*p*<0.001 vs LXA4 (10 nM) + agomiR-NC.

### SIRT1 participated in the process of LXA4 promoting fat browning

By constructing and transfecting the overexpression/silence plasmids of SIRT1, the participation of SIRT1 participated in the promotive effects of LXA4 on fat browning was proved. Transfection efficiency test validated that the plasmids of overexpressed SIRT1, shSIRT1-1 and shSIRT1-2 evidently changed the expression of SIRT1 in cells (*p*<0.01, figure 6A). Among them, the down-regulation of shSIRT1-2 exerted a more significant impact, so the shSIRT1-2 was employed for subsequent analyses.

**Figure 6.**
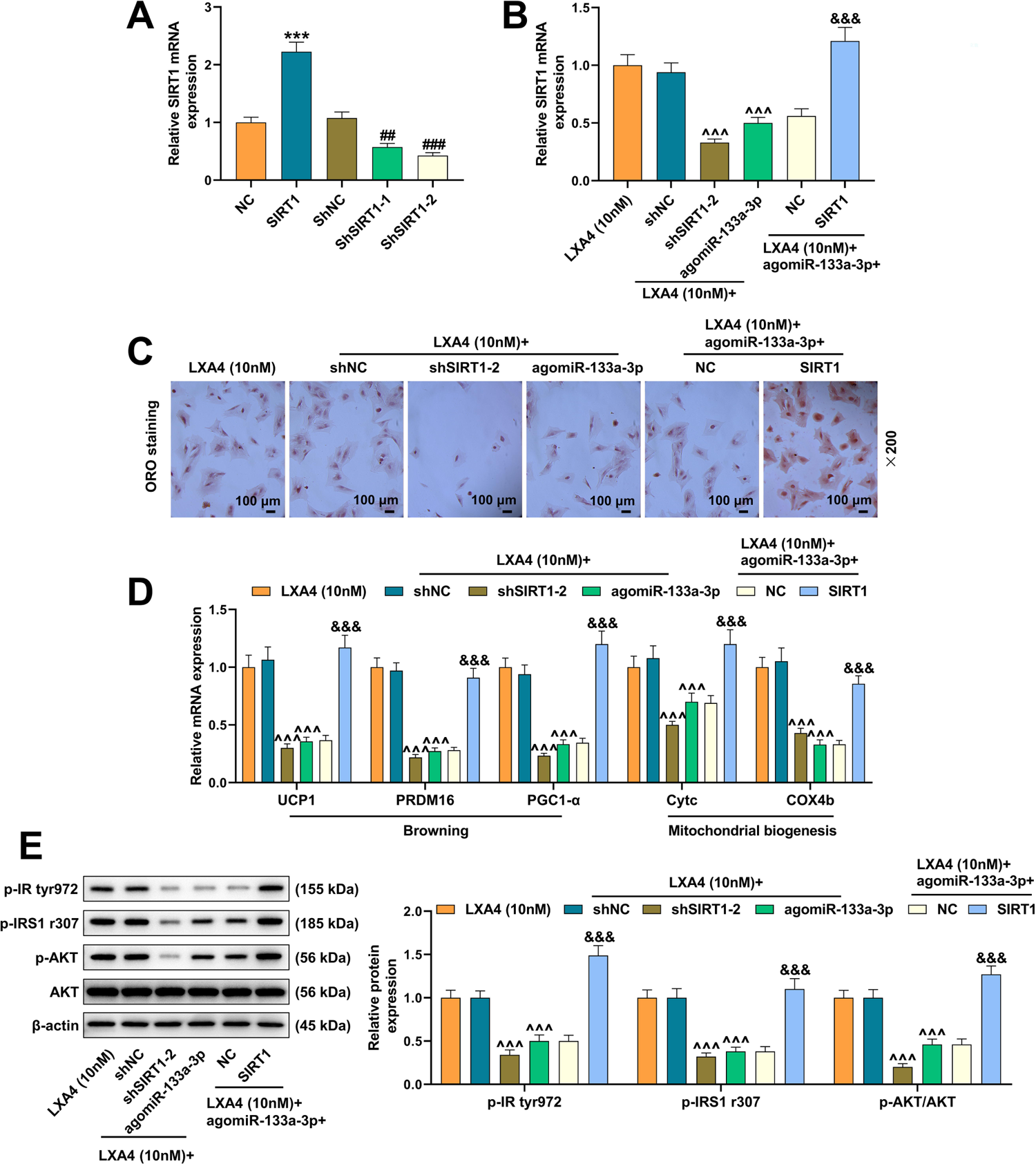
SIRT1 participated in the process of LXA4 promoting fat browning. (A) The transfection efficiency of overexpressed/silenced plasmid of Sirtuin1 (SIRT1) was measured by qRT-PCR. β-actin acted as the housekeeping gene. (B) The transfection efficiency of shSIRT1-2 and overexpressed SIRT1 in 3T3-L1 cells treated with LXA4 were tested by qRT-PCR. β-actin acted as the housekeeping gene. (C) The effects of silenced and overexpressed SIRT1 on lipid droplet formation in 3T3-L1 cells treated with LXA4 were observed by oil red O staining. The magnification was 200 times. (D) The effects of silenced and overexpressed SIRT1 on the expression of adipocyte browning-related genes in 3T3-L1 cells treated with LXA4 were tested by qRT-PCR. β-actin acted as the housekeeping gene. (E) The regulation of silenced and overexpressed SIRT1 on insulin receptor-AKT pathway in LXA4-treated 3T3-L1 cells was detected by western blot. β-actin acted as the housekeeping gene. All experiments were repeated three times to average. \*\*\**p*<0.001 vs NC; ##*p*<0.01, ###*p*<0.001 vs ShNC; ^^^*p*<0.001 vs LXA4 (10 nM) + ShNC; &&&*p*<0.001 vs LXA4 (10 nM) + agomiR-133a-3p + NC.

In the next rescue experiments, the promotive effect of LXA4 and the inhibitory effect of agomiR-133a-3p on SIRT1 expression as well as adipocyte lipid droplet accumulation, fat browning-related factors and insulin receptor-AKT pathway (*p*<0.001, figure 6C-E) were markedly overturned by the transfection of shSIRT1-2 and SIRT1 overexpression plasmid, respectively (*p*<0.001, figure 6B-E).

## Discussion

With the advancement of social economy and the changes in people’s lifestyles and living standards, obesity, accompanied by increased incidence year by year, has become a worldwide public health problem. As the two fat forms of mammals, WAT and BAT have obvious differences in the morphology of their constituent cells (Cinti, 2012, Frontini and Cinti, 2010): White adipocytes contain only one large cytoplasmic lipid droplet, while brown adipocytes are rich in oxygen supply and innervated by sympathetic nerves. They contain not only many small lipid droplets, but also numerous mitochondria in the cytoplasm. It is this structural difference that determines their functional difference. Promoting WAT browning and BAT activity through various physiological or pharmacological methods is currently a new strategy to combat obesity (Bonet et al., 2013, Calvani et al., 2014), which acted as a supplementary treatment for this study. In animal experiments, LXA4 successfully lessened the body weight and fat mass of food-induced obese mice. Further pathological analysis of WAT/BAT verified that LXA4 dwindled the volume of white adipocytes and promoted that of brown adipocytes, which is consistent with Elias’s conclusion, suggesting that LXA4 promotes the browning of white adipocytes.

Therefore, in the next *in vitro* study, we analyzed the regulation of LXA4 on the browning-related genes of adipocytes in differentiated 3T3-L1 cells. Studies demonstrated that UCP1 was activated during cold stimulation, and increased oxidation of blood sugar and free fatty acids after ingestion, maintaining a higher level of uncoupling in the BAT mitochondrial oxidative respiratory chain and generating more heat (Marlatt and Ravussin, 2017). PRDM16 and PGC1-[are markers of brown adipocyte maturation (Frontini and Cinti, 2010). The up-regulation of Cytc and COX4b reflects the activation of mitochondrial function (Medar and Marinkovic, 2020). Our results corroborated that the MDI inducer successfully activated the above genes, and the addition of LXA4 further up-regulated the browning-related gene expressions of adipocytes, denoting that LXA4 combined with MDI generated better effects than MDI alone.

Recently, the advances in genomics and bioinformatics have led to a growing recognition of the important role of non-coding RNA in human physiology and pathology. MiR-133a-3p, miR-27b-3p, miR-203, miR-503, and miR-429 can all regulate the browning process of white adipocytes (Kim et al., 2019, Petrek and Yu, 2019, Guo et al., 2019, Man et al., 2020, Ye et al., 2021). In the light of analyses on the regulation of expression and the verification of rescue test, we finally determined that LXA4 promotes the browning of 3T3-L1 adipocytes by regulating miR-133a-3p. At first, miR-133a-3p was confirmed as a tumor suppressor gene in colorectal, ovarian, and bladder cancers (Liang et al., 2018, Yu et al., 2019). With the deepening of research, scientists have gradually unearthed the role of miR-133a-3p in other diseases. For example, Li proposed that miR-133a-3p is involved in the apoptosis of cardiomyocytes, acting as a potential therapeutic target for ischemic cardiomyopathy (Li et al., 2018). Based on the functional mechanism of miRNA-mRNA, we found the downstream target gene of miR-133a-3p, SIRT1, which belongs to the silent information regulator family and plays a pivotal role in the processes of glucose and lipid metabolism, as well as cell differentiation, inflammation and apoptosis. A number of studies have certified (Zhou et al., 2019, Li et al., 2020) that the SIRT1 pathway mediates adipocyte browning and suppresses fat accumulation. These arguments fully explain our experimental results.

Obesity is one of the main causes of insulin resistance, and lipolysis releases considerable free fatty acids. Excessive free fatty acid content will cause lipotoxicity, reduce the sensitivity of muscle, fat, and liver to insulin, and at the same time affect the function of pancreatic β-cells to secrete insulin (Capurso and Capurso, 2012). These phenomena were also observed in our experiments. HFD caused a large accumulation of glucose in mice and reduced insulin sensitivity. Therefore, in terms of molecular mechanism selection, we analyzed insulin-related changes in the insulin receptor-AKT pathway, manifesting that LXA4 activated p-IR tyr972, p-IRS1 r307 and p-AKT in this pathway. IRS1 is an important signal protein at the post-receptor level in the insulin signal transduction pathway, distributed in all peripheral tissues such as skeletal muscle, fat, liver, etc., where insulin acts (Kubota et al., 2017). Tyr972 near the membrane domain is necessary for the insulin receptor β subunit to bind to IRS1, playing a key role in the transmission of insulin signals (Blesson et al., 2014). After binding to the insulin receptor, phosphorylated IRS1 can activate the p110 subunit of PI3K and the downstream signaling molecule Akt through an enzymatic reaction (Aoki and Terauchi, 2018).

In line with the above evidence, this study confirmed that LXA4 activates the insulin receptor-AKT pathway via regulating the miR-133a-3p/SIRT1 axis, ultimately promoting the browning of white adipocytes and alleviating insulin resistance. Moreover, this study supplemented the specific mechanism via which LXA4 promoted adipocyte browning, and provided a new reference for the treatment of clinical obesity. Of course, there is still room for improvement in our research. Validating the expression changes of the insulin receptor-AKT pathway through animal models and exploring the node changes between SIRT1 and pathways are the directions of our follow-up research.

## Acknowledgements

Not applicable.

## Data availability

The analyzed data sets generated during the study are available from the corresponding author on reasonable request.

